# Effect of obesity-linked *FTO* rs9939609 variant on physical activity and dietary patterns in physically active men and women

**DOI:** 10.1101/091728

**Authors:** Nathan R. West, James Dorling, Alice E. Thackray, Samantha E. Decombel, David J. Stensel, Stuart J. Grice

## Abstract

**Objective:** To examine physical activity and dietary patterns in physically active individuals carrying different risk variants of the fat mass and obesity-associated gene (*FTO*) rs9939609 single nucleotide polymorphism (SNP).

**Methods:** A total of 528 white men and women (mean (SD): 34.9(9.5) years, 26.6(4.3) kg·m^-2^) were genotyped for *FTO* rs9939609 SNP. Sex, age and anthropometric measurements (stature, body mass, waist circumference) were self-reported using an online questionnaire, and body mass index and waist-to-height ratio were calculated. Physical activity level and eating behaviour were assessed using the International Physical Activity Questionnaire and Three-Factor Eating Questionnaire (TFEQ), respectively.

**Results:** Body mass, body mass index, waist circumference and waist-to-height ratio were not significantly different between individuals carrying different *FTO* rs9939609 risk variants (all P≥0.66). The cohort was physically active (4516(3043) total MET min·week^-1^), although risk allele carriers (AT/AA) reported higher total physical activity (effect size = 0.22, p=0.03), and homozygous risk allele carriers (AA) displayed higher TFEQ cognitive restraint (effect size = 0.33, p=0.03) compared with non-risk allele carriers (TT).

**Conclusions:** Obesity-related parameters were not different in physically active individuals carrying different risk variants of *FTO* rs9939609, but higher physical activity and cognitive restraint in risk allele carriers may reduce genetic predisposition to weight gain.

## Introduction

Obesity is a major risk factor for several chronic diseases and represents a major health and economic burden on society (1). The aetiology of obesity is multifactorial, and is influenced by complex interactions between environmental, lifestyle and genetic factors (2). Consequently, it is important to understand the interplay between these factors when designing strategies targeting the prevention of obesity.

The fat mass and obesity-associated gene (*FTO*) was the first common variant identified by genome wide association studies that influences obesity risk (3). Single nucleotide polymorphisms (SNPs) in intron 1 of *FTO* have been associated consistently with obesity risk across different ages and populations (4–9). At *FTO* rs9939609 SNP, individuals homozygous for the risk allele (AA) weigh 3 kg more, and have a 1.7-fold higher risk of obesity than those who do not carry a risk allele (TT) (3). Studies have examined the effect of *FTO* variants on regulators of energy homeostasis to elucidate the mechanisms influencing *FTO*-mediated obesity risk. In this respect, evidence suggests that *FTO* may play a central role in the regulation of food intake (10,11). This is supported by studies demonstrating that individuals homozygous for the risk allele exhibit reduced satiety, poor food choices and increased energy consumption (12–14). Conversely, *FTO* obesity SNPs have not been related to energy expenditure, with evidence suggesting that those carrying the risk allele do not show reduced basal metabolic rate (15) or physical activity levels (7,16,17).

Although the association between *FTO* and obesity risk is well established, lifestyle may modulate obesity risk related to *FTO.* Several studies have suggested that FTO-mediated body fatness may be attenuated in physically active individuals (18–20). Indeed, a meta-analysis concluded that higher physical activity levels attenuate the influence of *FTO* variation on obesity risk by 30% (21), and exercise interventions have demonstrated efficacy in promoting weight loss in *FTO* risk allele carriers (22,23). However, it is not well understood how the body mass index (BMI)-increasing influence of *FTO* is attenuated in physically active individuals. An improved understanding of the differences in dietary and physical activity patterns in variants of *FTO* rs9939609 SNP amongst a physically active cohort may provide a greater insight into the behaviours that offset FTO-mediated obesity. Therefore, the aim of this study was to examine physical activity and dietary habits in a sample of physically active men and women carrying different risk variants of *FTO* rs9939609 SNP.

## Methods

### Participants

With the approval of Loughborough University’s Ethical Advisory Committee, 708 men and women were recruited between March to November 2015 to participate in this study. Participants were recruited through FitnessGenes Ltd. (Bicester, Oxfordshire, UK - previously MuscleGenes Ltd.) and were predominantly based in European countries or the United States. Full informed consent, via an online consent form, was obtained from participants before the study commenced. Participants classified themselves into one of fifteen ethnic groups including three categories of White descent (British, Irish, other White background). Data presented in this study is from participants classifying themselves as White. Of the 708 recruited, 1 had missing genotype data, 17 had missing demographic or anthropometric data and 162 did not classify themselves as White. Therefore, results are presented for 528 participants (421 males, 107 females). Participant characteristics are presented in Table 1.

**Table 1.** Participant characteristics.

|  | **Total**<br>**(n = 528)** | **Female**<br>**(n = 107)** | **Male**<br>**(n = 421)** |
| --- | --- | --- | --- |
| **Age and anthropometric characteristics** |  |  |  |
| Age (years) | 34.9 (9.5) | 36.8 (9.1) | 34.5 (9.6)* |
| Stature (cm) | 176.5 (9.4) | 164.9 (7.1) | 179.5 (7.4)* |
| Body mass (kg) | 83.4 (16.5) | 69.0 (13.9) | 87.0 (15.0)* |
| Body mass index (kg·m<sup>-2</sup>) | 26.6 (4.3) | 25.4 (4.8) | 27.0 (4.0)* |
| Waist circumference (cm) | 84.6 (10.4) | 78.2 (11.8) | 86.2 (9.4)* |
| Waist-to-height ratio | 0.48 (0.06) | 0.48 (0.07) | 0.48 (0.05) |
| Central obesity<sup>a</sup> | 37 (7%) | 21 (20%) | 16 (4%)** |
| **Physical activity levels<sup>b</sup>** |  |  |  |
| Vigorous MET min·week<sup>-1</sup> | 2629 (1908) | 2511 (1912) | 2659 (1908) |
| Moderate MET min·week<sup>-1</sup> | 737 (1026) | 716 (874) | 743 (1062) |
| Walking MET min·week<sup>-1</sup> | 1150 (1269) | 1284 (1374) | 1115 (1241) |
| Total MET min·week<sup>-1</sup> | 4516 (3043) | 4511 (3163) | 4518 (3017) |
| **Eating behaviour** |  |  |  |
| Cognitive restraint score | 12 (4) | 12 (4) | 12 (4) |
| Disinhibition score | 6 (4) | 7 (4) | 6 (3)* |
| Hunger score | 5 (4) | 6 (4) | 5 (4) |
| ***FTO* rs99396909 genotype** |  |  |  |
| AA | 95 (18%) | 18 (17%) | 77 (18%) |
| AT | 236 (45%) | 40 (37%) | 196 (47%) |
| TT | 197 (37%) | 49 (46%) | 148 (35%) |

Values for age, anthropometric characteristics, physical activity levels and eating behaviour are mean (SD) and were analysed between sex using linear mixed models. Values for central obesity and *FTO* rs99396909 genotype are frequency (%) and were analysed between sex using Chi-square tests.
<sup>a</sup> Central obesity was defined as a waist circumference > 88 cm for women and > 102 cm for men
<sup>b</sup> Complete physical activity data available for *n* = 408 (female *n* = 83, male *n* = 325)
\* Significant difference between females and males (linear mixed model *P* < 0.05)
\*\* Significant difference between females and males (Chi-square test *P* < 0.05)

### Genotype analysis

Participant DNA was obtained from saliva, which was collected via an Oragene DNA self-collection kit (DNA Genotek Inc., Ottawa, ON, Canada) sent from and returned to FitnessGenes Ltd. by post. DNA was extracted by LGC Genomics (Hertfordshire, UK) and genotyped using a KASP™ genotyping assay for *FTO* rs9939609 SNP. Genotype frequency of *FTO* rs9939609 SNP was assessed using a goodness-of-fit chi-square test and did not deviate from Hardy-Weinberg equilibrium (AA = 95, AT = 236, TT = 197; P = 0.10).

### Collection of demographic and anthropometric data

Participants self-reported their sex, age, stature, body mass, waist circumference, country of birth and ethnicity using an online questionnaire. Sex, age and ethnicity were confirmed by cross-referencing against customer information held by FitnessGenes Ltd. Participant stature, body mass and waist circumference were used to calculate BMI and waist-to-height ratio (Table 1). Central obesity was calculated as the percentage of participants that exceeded previously defined thresholds for waist circumference (females > 88 cm, males > 102 cm) (24).

### Physical activity levels

Physical activity levels were measured using the validated short format of the International Physical Activity Questionnaire (IPAQ) (25). The IPAQ assesses the frequency and duration of walking, moderate- and vigorous-intensity physical activities performed in bouts lasting 10 minutes or more during the previous seven days. Total physical activity (MET minutes per week) was estimated by multiplying the number of minutes reported in each activity level by the specific MET score for that activity (walking: 3.3, moderate-intensity: 4.0, vigorous-intensity: 8.0 METs), and participants were classified in one of three physical activity levels: low, moderate or high (http://www.ipaq.ki.se).

### Eating behaviour

Eating behaviour was assessed using the validated 51-item Three-Factor Eating Questionnaire (TFEQ) to measure dietary restraint (21 items, Cronbach’s α 0.788), disinhibition (16 items, Cronbach’s α 0.745) and hunger (14 items, Cronbach’s α 0.761) (26). All TFEQ items were coded with either 0 or 1 point and summed within each domain. Higher scores within each domain were indicative of higher restrained eating, disinhibited eating or a predisposition to hunger.

### Statistical analyses

Data were analysed using the IBM SPSS Statistics Software for Windows Version 21 (IBM, New York). Between-sex differences in participant characteristics were examined using linear mixed models with one fixed factor (sex). Linear mixed models, adjusted for age and sex, were used to examine between-genotype differences in obesity-related parameters, physical activity levels and eating behaviour with one fixed factor *(FTO* rs99396909 genotype). Between-genotype differences in all outcome measures were analysed using: 1) genotype model (AA vs. AT vs. TT); 2) dominant model (risk allele carriers (AA/AT) vs. homozygous non-risk genotype (TT)); and 3) recessive model (non-risk allele carriers (AT/TT) vs. homozygous risk genotype (AA)). All linear mixed models included a random effect for each participant. Where significant main effects were found in the genotype model, post hoc analysis was performed using Holm-Bonferroni correction for multiple comparisons. Differences in categorical variables (central obesity, *FTO* genotype frequency) between sex and/or *FTO* genotype groups were analysed using Chi-square tests. Effect sizes are used to supplement important findings. An effect size of 0.2 was considered the minimum important difference in all outcome measures, 0.5 moderate and 0.8 large (27). Continuous variables are presented as mean (SD) and categorical variables as frequency (%). Statistical significance was accepted as P < 0.05.

## Results

### Participant characteristics

On average, participants reported engaging in a total of 4516 (3043) MET minutes of activity per week and the majority of participants were classified in the high physical activity category (high 75.2%, moderate 19.4%, low 5.4%), confirming the cohort were physically active (Table 1). Participants reported the following reasons for engaging in physical activity: muscle building training *n* = 276 (52%); fat loss training *n* = 168 (32%); strength training *n* = 40 (8%); power training *n* = 20 (4%); endurance training *n* = 17 (3%); no response *n* = 7 (1%).

Females were significantly older (ES = 0.24, P = 0.02), had a higher frequency of central obesity (odds ratio = 6.31, P < 0.001) and exhibited higher TFEQ disinhibition scores (ES = 0.42, P < 0.001) compared with males (Table 1). Body mass, BMI and waist circumference were significantly lower in females than males (all ES ≥ 0.38, P ≤ 0.001), but waist-to-height ratio was similar between the sexes (ES = 0.09, P = 0.40) (Table 1). No significant differences were seen between the sexes for any measure of physical activity (all ES ≤ 0.13, P ≥ 0.28) or *FTO* genotype frequency (P = 0.12) (Table 1).

### FTO rs9939609 genotype and obesity-related parameters

Obesity-related parameters in men and women carrying different risk variants of *FTO* rs9939609 SNP are displayed in Figure 1 and Table 2. Linear mixed models revealed no significant differences in body mass, BMI, waist circumference or waist-to-height ratio across *FTO* rs9939609 genotype groups (all P ≥ 0.66). No significant differences were seen in obesity-related parameters when analysed in dominant (all ES ≤ 0.13, P ≥ 0.37) or recessive (all ES ≤ 0.08, P ≥ 0.68) allele models.

**Figure 1.**
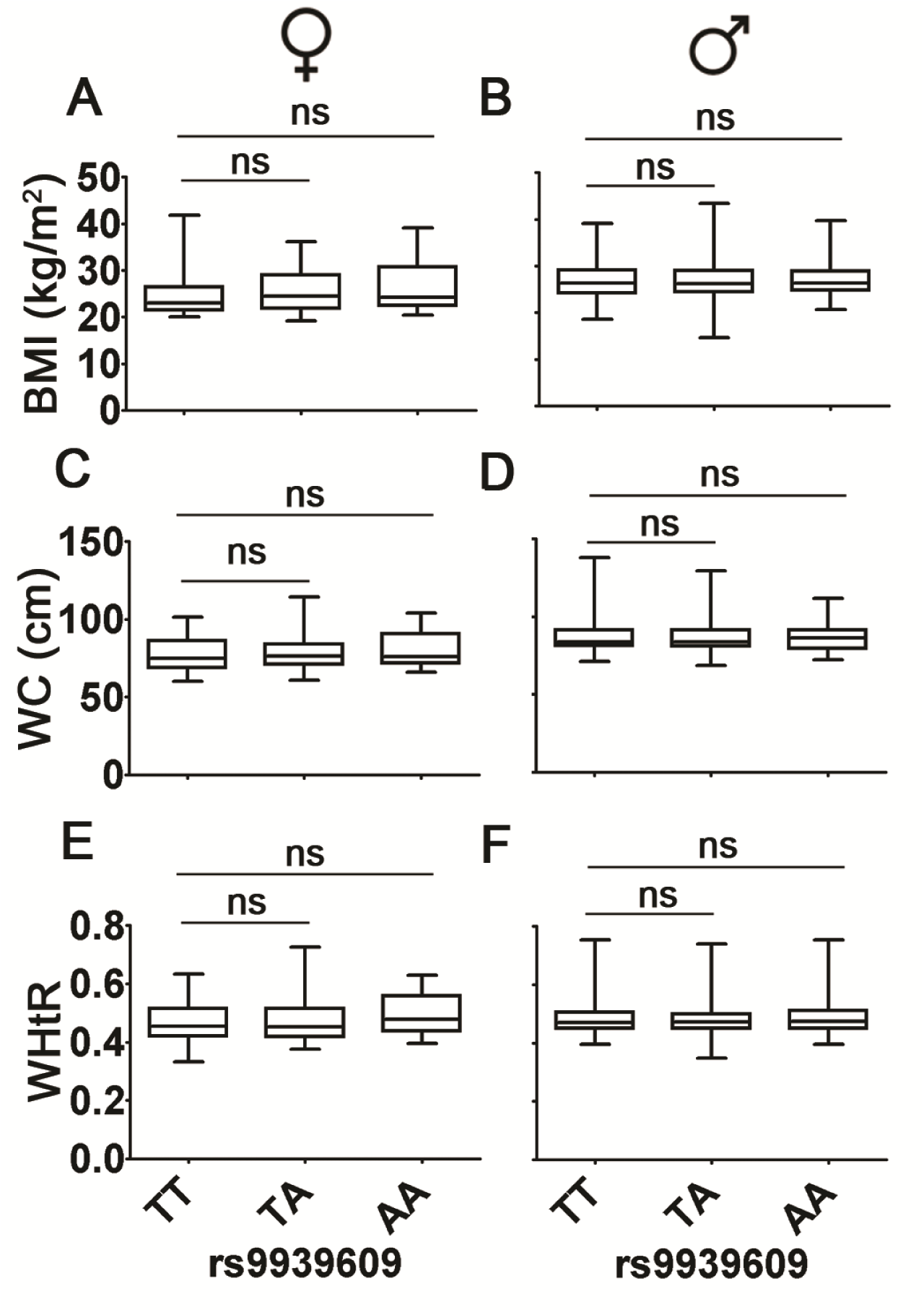
Box plots and analysis of the effect of *FTO* rs9939609 gene variant on obesity-related parameters. Panel A, B: body mass index (BMI); C, D: waist circumference (WC); E, F: waist-to-height ratio (WHtR). Panel A, C, E: female; B, D, F: male. Females *n* = 107 (AA = 18, AT = 40, TT = 49). Males *n* = 421 (AA = 77, AT = 196, TT = 148). Line within box represents the median while the lower and upper box lines represent the interquartile range (1^st^ and 3^rd^ quartile). The whiskers represent the minimum and maximum observations. Linear mixed models, adjusted for age and sex, were used to examine between genotype differences. No significant differences were seen in obesity-related parameters between *FTO* rs9939609 genotypes.

**Table 2.** Obesity-related parameters in men and women carrying different risk variants of the *FTO* rs99396909 single nucleotide polymorphism.

| Characteristic | *FTO* rs99396909 genotype |  |  |  |  | Model 1<sup>a</sup>:<br>Genotype<br>P value | Model 2<sup>b</sup>:<br>Dominant<br>P value<br>(ES/OR) | Model 3<sup>c</sup>:<br>Recessive<br>P value<br>(ES/OR) |
| --- | --- | --- | --- | --- | --- | --- | --- | --- |
|  | AA | AT | TT | AA/AT | AT/TT |  |  |  |
| Body mass (kg) | 83.7 (14.7) | 84.4 (16.5) | 82.0 (17.2) | 84.2 (16.0) | 83.3 (16.8) | 0.69 | 0.44 (0.13) | 0.93 (0.02) |
| Body mass index (kg·m<sup>-2</sup>) | 26.8 (3.9) | 26.8 (4.3) | 26.4 (4.3) | 26.8 (4.2) | 26.6 (4.3) | 0.66 | 0.37 (0.09) | 0.79 (0.05) |
| Waist circumference (cm) | 85.2 (9.4) | 84.9 (10.4) | 84.1 (11.0) | 85.0 (10.1) | 84.5 (10.7) | 0.87 | 0.60 (0.09) | 0.88 (0.07) |
| Waist-to-height ratio | 0.48 (0.05) | 0.48 (0.06) | 0.48 (0.06) | 0.48 (0.05) | 0.48 (0.06) | 0.85 | 0.61 (0.03) | 0.68 (0.08) |
| Central obesity<sup>d</sup> | 6 (7%) | 16 (7%) | 15 (8%) | 22 (7%) | 31 (7%) | 0.92 | 0.70 (0.87) | 0.80 (1.12) |

Values for body mass, body mass index, waist circumference and waist-to-height ratio represent mean (SD) and were analysed using linear mixed models adjusted for age and sex. Values for central obesity represent frequency (%) and were analysed using Chi-square tests. ES, effect size (body mass, body mass index, waist circumference, waist-to-height ratio); OR, odds ratio (central obesity).
<sup>a</sup> Model 1: genotype model (AA vs. AT vs. TT)
<sup>b</sup> Model 2: dominant model (AA/AT vs. TT)
<sup>c</sup> Model 3: recessive model (AT/TT vs. AA)
<sup>d</sup> Central obesity was defined as a waist circumference > 88 cm for women and > 102 cm for men

### FTO rs9939609 genotype and physical activity levels

Complete physical activity data were available for 408 participants (AA *n* = 74, AT *n* = 177, TT *n* = 157; female *n* = 83, male *n* = 325). Physical activity levels in men and women carrying different risk variants of *FTO* rs9939609 SNP are displayed in Table 3. Linear mixed models revealed no significant differences in physical activity levels across *FTO* rs9939609 genotype groups (all P ≥ 0.10). The dominant allele model revealed total MET minutes per week (ES = 0.22, P = 0.03) and a tendency for vigorous MET minutes per week (ES = 0.19, P = 0.08) to be higher in A allele carriers (AA/AT) than non-risk allele carriers (TT). No significant differences in moderate or walking MET minutes per week were identified in the dominant allele model (AA/AT vs. TT) (all ES ≤ 0.13, P ≥ 0.17). The recessive allele model (AT/TT vs. AA) revealed no significant differences in physical activity levels between *FTO* genotype groups (all ES ≤ 0.11, P ≥ 0.28).

**Table 3.** Physical activity levels in men and women carrying different risk variants of the FTO rs99396909 single nucleotide polymorphism.

| Characteristic | *FTO* rs99396909 genotype |  |  |  |  | Model 1<sup>a</sup>:<br>Genotype<br>P value | Model 2<sup>b</sup>:<br>Dominant<br>P value (ES) | Model 3<sup>c</sup>:<br>Recessive<br>P value (ES) |
| --- | --- | --- | --- | --- | --- | --- | --- | --- |
|  | AA (*n* = 74) | AT (*n* = 177) | TT (*n* = 157) | AA/AT | AT/TT |  |  |  |
| Vigorous MET min·week<sup>-1</sup> | 2808 (1966) | 2751 (1903) | 2407 (1876) | 2768 (1918) | 2589 (1895) | 0.19 | 0.08 (0.19) | 0.29 (0.11) |
| Moderate MET min·week<sup>-1</sup> | 726 (1069) | 816 (1127) | 654 (874) | 789 (1109) | 740 (1018) | 0.41 | 0.21 (0.13) | >0.99 (0.01) |
| Walking MET min·week<sup>-1</sup> | 1240 (1242) | 1207 (1345) | 1043 (1192) | 1217 (1313) | 1130 (1276) | 0.34 | 0.17 (0.14) | 0.32 (0.09) |
| Total MET min·week<sup>-1</sup> | 4774 (3125) | 4774 (3193) | 4104 (2795) | 4774 (3167) | 4459 (3027) | 0.10 | 0.03 (0.22)** | 0.28 (0.10) |

Values are mean (SD) for *n* = 408. Comparisons were made using linear mixed models adjusted for age and sex. ES, effect size.
<sup>a</sup> Model 1: genotype model (AA vs. AT vs. TT)
<sup>b</sup> Model 2: dominant model (AA/AT vs. TT)
<sup>c</sup> Model 3: recessive model (AT/TT vs. AA)
\*\* Significant difference between AA/AT and TT *FTO* rs99396909 genotype (linear mixed model *P* < 0.05)

### FTO rs9939609 genotype and eating behaviours

Eating behaviours in men and women carrying different risk variants of *FTO* rs9939609 SNP are displayed in Table 4. Linear mixed models identified a significant main effect for cognitive restraint score across *FTO* genotype groups (P = 0.03). Post hoc analysis of between-group differences revealed that the cognitive restraint score was higher in homozygous A allele carriers than AT (ES = 0.25, P = 0.07) and TT (ES = 0.33, P = 0.03) genotypes; AT and TT genotypes were similar (ES = 0.06, P = 0.49). No differences in disinhibition or hunger scores were seen across *FTO* genotype groups (both P ≥ 0.38).

**Table 4.** Eating behaviour in men and women carrying different risk variants of the *FTO* rs99396909 single nucleotide polymorphism.

| Characteristic | *FTO* rs99396909 genotype |  |  |  |  | Model 1<sup>a</sup>:<br>Genotype<br>P value | Model 2<sup>b</sup>:<br>Dominant<br>P value (ES) | Model 3<sup>c</sup>:<br>Recessive<br>P value (ES) |
| --- | --- | --- | --- | --- | --- | --- | --- | --- |
|  | AA (*n* = 95) | AT (*n* = 236) | TT (*n* = 197) | AA/AT | AT/TT |  |  |  |
| Cognitive restraint score | 13 (4) | 12 (4) | 11 (4) | 12 (4) | 11 (4) | 0.03* | 0.12 (0.14) | 0.01 (0.28)*** |
| Disinhibition score | 6 (3) | 7 (4) | 6 (4) | 6 (4) | 6 (4) | 0.38 | 0.32 (0.07) | 0.60 (0.08) |
| Hunger score | 5 (3) | 5 (4) | 5 (4) | 5 (4) | 5 (4) | 0.92 | 0.99 (0.02) | 0.69 (0.01) |

Values are mean (SD) for *n* = 528. Comparisons were made using linear mixed models adjusted for age and sex. ES, effect size.
<sup>a</sup> Model 1: genotype model (AA vs. AT vs. TT)
<sup>b</sup> Model 2: dominant model (AA/AT vs. TT)
<sup>c</sup> Model 3: recessive model (AT/TT vs. AA)
\* Significant difference between AA and TT *FTO* rs99396909 genotype (linear mixed model $P < 0.05$ after Holm-Bonferroni correction)
\*\*\* Significant difference between AT/TT and AA *FTO* rs99396909 genotype (linear mixed model $P < 0.05$ )

The dominant allele model (AA/AT vs. TT) revealed no significant differences in cognitive restraint, disinhibition or hunger scores between *FTO* genotype groups (all ES ≤ 0.14, P ≥ 0.12). The recessive allele model revealed that the cognitive restraint score was higher in homozygous A allele carriers (AA) than T allele carriers (AT/TT) (ES = 0.28, P = 0.01). No significant differences in disinhibition or hunger scores were identified in the recessive allele model (all ES ≤ 0.08, P ≥ 0.60).

## Discussion

The primary finding from the present study was that obesity-related parameters were not different in physically active individuals carrying different risk variants of *FTO* rs9939609 SNP. Furthermore, *FTO* rs9939609 A allele carriers exhibited greater physical activity levels and cognitive restraint than non-risk allele carriers. This suggests that physical activity and diet behaviours may reduce genetic predisposition to weight gain and obesity within a physically active cohort.

The consensus of evidence suggests that *FTO* risk alleles are associated with elevated body weight across different ages and populations, with each minor risk allele increasing BMI and obesity risk by 0.25–0.39 kg·m^-2^ and 1.18–1.27 fold, respectively (9). However, the most extensively studied rs9939609 SNP had no influence on obesity-related parameters in the current study. Several studies have demonstrated using self-reported questionnaires (28,29) and objective physical activity devices (19,30) that obesity-related traits of *FTO* are attenuated in individuals with higher physical activity levels. Therefore, the high physical activity levels amongst the current cohort may have negated any differences in obesity-related parameters between *FTO* rs9939609 genotypes. Intriguingly, despite being a highly active group when taken as a whole, the current study demonstrated that individuals with the A allele reported higher total and vigorous physical activity levels compared with TTs. It is possible, therefore, that this difference in physical activity patterns between *FTO* genotypes may further offset the adiposity-increasing effect of *FTO* within this cohort. Nonetheless, this elevation in activity was small and differences in physical activity levels between *FTO* genotypes conflicts with previous studies suggesting that *FTO* genotype does not impact on physical activity levels (7,28). Contradictions between the current study and other evidence may be due to the high levels of physical activity of individuals within the current study. Although Berentzen et al. (16) reported no effect of *FTO* variant on physical activity in individuals classified by the authors as highly active (more than 4 h moderate physical activity a week), there is a lack of studies that have examined the effect of *FTO* genotype on activity levels in cohorts with even greater levels of physical activity. Consequently, further studies are required to examine *FTO* differences in physical activity amongst individuals with varying activity status. Moreover, as much of the data on *FTO*-mediated differences in physical activity levels are reliant on self-reported questionnaire data, additional scientific enquiry using objective physical activity measures such as accelerometers is required.

It has been postulated that *FTO*-mediated predisposition to weight gain and obesity may be modified by dietary behaviours. Research suggests that *FTO* may play a significant role in the regulation of satiety and food intake (10,11). Furthermore, it has been reported in previous observational studies that the risk variant of *FTO* is associated with higher cognitive restraint, disinhibition and hunger which may be indicative of poorer eating behaviours (31,32). The present study extends these findings by demonstrating that physically active individuals homozygous for *FTO* rs9939609 risk allele display higher cognitive restraint but similar disinhibition and hunger scores than non-risk allele carriers. These small differences in eating behaviour may attenuate FTO-mediated obesity risk in physically active individuals. The potential benefit of cognitive restraint on the propensity to body weight changes is highlighted by studies demonstrating a negative association between cognitive restraint and markers of adiposity (e.g., body mass, BMI) (33,34). However, such associations are not reported universally (35), and cognitive restraint tends to be higher in overweight compared with healthy weight individuals (36). Additionally, disinhibition has been proposed as a stronger predictor of BMI (37). Nevertheless, increased dietary restraint during weight loss has been identified as a significant predictor of successful weight loss maintenance (38). Therefore, it is possible that the higher cognitive restraint observed in *FTO* risk allele carriers in the present study may offset *FTO*-mediated obesity risk in physically active individuals, but further work is required to confirm this chronically.

The current study is not without limitations. First, the anthropometric, physical activity and eating behaviours data were self-measured and self-reported using an online survey, which may have led to problems with participant measurement error, recall and bias. Additional work is, therefore, required using more direct and objective measures. Second, one of the primary measures of body fatness in the present study was BMI. Despite its frequent use in large-scale population studies, BMI does not take into account differences in fat and fat-free mass such as muscle and bone (39). Given the current cohort reported high levels of total activity with the majority indicating their primary fitness goal was muscle building (52%), it is possible that variations in body mass (and hence BMI) could be largely due to higher muscle mass. This may explain why the average BMI amongst individuals in the current data set falls within the overweight category. Thus, future studies using more direct measures of body adiposity are required to replicate the current study’s findings.

## Conclusion

Within a physically active cohort, risk allele carriers of *FTO* rs9939609 exhibited greater physical activity levels and cognitive restraint than non-risk allele carriers, despite demonstrating similar adiposity-related measures. Consequently, physical activity and cognitive restraint may lower *FTO*-mediated susceptibility to weight gain and obesity within a physically active cohort. Future studies with repeated and objective measurements are required to further investigate physical activity and dietary behaviours that underscore the effects of *FTO* on obesity risk.

## Acknowledgements

The authors thank the volunteers for their participation in this study.

